# A 512-ch dual-mode microchip for simultaneous measurements of electrophysiological and neurochemical activities

**DOI:** 10.1101/2022.05.18.492473

**Authors:** Geoffrey Mulberry, Kevin A. White, Brian N. Kim

## Abstract

In the study of the brain, large and high-density microelectrode arrays have been widely used to study the behavior of neurotransmission. CMOS technology has facilitated these devices by enabling the integration of high-performance amplifiers directly on-chip. Usually, these large arrays measure only the voltage spikes resulting from action potentials traveling along firing neuronal cells. However, at synapses, communication between neurons occurs by the release of neurotransmitters, which cannot be measured on typical CMOS electrophysiology devices. Development of electrochemical amplifiers has resulted in the measurement of neurotransmitter exocytosis down to the level of a single vesicle. To effectively monitor the complete picture of neurotransmission, measurement of both action potentials and neurotransmitter activity is needed. Current efforts have not resulted in a device that is capable of the simultaneous measurement of action potential and neurotransmitter release at the same spatiotemporal resolution needed for a comprehensive study of neurotransmission. In this paper, we present a true dual-mode CMOS device that fully integrates 256-ch electrophysiology amplifiers and 256-ch electrochemical amplifiers, along with an on-chip 512 electrode microelectrode array capable of simultaneous measurement from all 512 channels. Electrochemical amplifiers achieve a noise level of 4.51 pA_RMS_ with a bandwidth of 10.3 kHz. Electrophysiology amplifiers exhibit 24.9 μV_RMS_ noise in the action potential band and a total usable bandwidth from 0.2 Hz to 10 kHz.

## I. Introduction

**T**he detailed study of synaptic function is of paramount importance for a better understanding of neurodegenerative diseases. Many diseases involving motor function, sensory, and cognitive impairments such as Parkinson’s Disease, Huntington’s Disease, and Alzheimer’s Disease are speculated to be caused by dysfunction of the synapses [1]–[5]. Alzheimer’s has been shown to develop in patients with an abnormal buildup and accumulation of amyloid-beta and tau oligomer proteins, which have been shown to harm the communication between synapses as well as neuronal loss [1], [6], [7]. Parkinson’s and another neurodegenerative disease, Lewy Body Dementia, are thought to develop from a similar underlying process through the buildup of alpha-synuclein and/or tau oligomers [8]–[10]. This degradation in synaptic function is common to all of these neurodegenerative diseases and is even noticed in many preclinical cases, suggesting the need for early detection of degradation [4], [11], [12].

There are primarily two types of electrical measurement used when studying neuronal activity: electrophysiology and amperometry. Electrophysiology is a broader term but within the study of the brain, it generally refers to measurements of voltage, particularly the measurement of Action Potentials (AP) and local field potential (LFP). An AP is also known as a “spike” or a “nerve impulse” and is caused when a neuron “fires” and causes a depolarization and repolarization to travel along the neuron’s axon. Amperometry involves the measurement of electrical current by fixing the potential of an electrode held near an active cell and measuring the current produced by the oxidation of neurotransmitters with a transimpedance amplifier (TIA). The process of secretion of neurotransmitters is known as exocytosis and is heavily studied using amperometry. Traditionally, researchers in a typical study perform either amperometry measurements or electrophysiology measurements, seldom both simultaneously. Additionally, these measurements are typically low throughput, involving in many cases only single electrodes.

Recent developments in the field have begun to use CMOS technology to integrate large numbers of amplifiers and microelectrode arrays (MEA) onto a single chip for high throughput electrochemical measurements [13], [14]. Presently, there is no current technology that exists at the microscopic scales to measure both modes of electrical activity involved in neurotransmission. Without this dual-mode capability, it is difficult to accurately monitor the dynamics of neuronal degradation. Most current studies are relying on measurements using microelectrode arrays to measure action potentials as they travel along axons to synapses. But these action potentials themselves are caused by secretions of neurotransmitters near the synapses of the neurons [15]–[17]. Thus, measuring action potentials alone, as is commonly done, is an indirect measurement which is not seeing the complete picture. If a complete understanding of the normal and dysfunctional neurotransmission is to be developed, a technology that can measure neurotransmitter release and action potentials simultaneously and at a small enough scale to actually capture relevant information at a statistically significant quantity. The integrated circuit presented in this work is the first of its kind being presented to solve these problems.

Many CMOS chips have been presented that focus on providing a high-density MEA capable of performing electrophysiology [13], [18]–[21]. Unfortunately, since they are performing electrophysiology, typically using a transconductance amplifier (TCA), they are incapable of measuring neurotransmitter release. However, there has been at least one recently-reported high-density CMOS MEA device that includes a small number of electrochemical amplifiers to measure neurotransmitter release in addition to typical electrophysiology measurements [22]. This device has a large number of electrodes (59,760), but can only measure from a selected portion of them (2048) at a time. Additionally, it only contains 28 amplifiers for measuring neurotransmitter release which have a noise performance on the order of 100s of pA_RMS_. The low amplifier count and poor noise performance make this device incapable of measuring neurotransmitter release at a similar spatiotemporal resolution as its electrophysiology capability.

In this paper, we present a new dual-mode chip that can simultaneously measure action potential propagations and neurochemical secretions at the synapses. Neurons can be measured using the dual-mode chip as demonstrated in Fig. 1a, which shows a cross-sectional view of a pair of neurons sitting above the surface of the device. By integrating both neurochemical and action potential amplifiers onto a single chip, presynaptic action potentials, neurotransmitter release at the synapse, and postsynaptic action potentials can be observed simultaneously. If a large quantity of both modalities of amplifier are arranged into a large array, then the flow of action potentials and neurotransmitters across complex neuronal networks can be studied (Fig. 1b). The dual-mode device presented in this paper is capable of measuring both neurotransmitter release and action potentials from 512 on-chip electrodes simultaneously and at the same spatiotemporal resolution. This device enables the study of the dynamic relationship between neurotransmission and action potential propagation using a single CMOS chip and MEA. Expanding on our previous work with high-density CMOS electrochemical MEAs [14], [23], we also add additional functionality to enable on-chip fast-scan cyclic voltammetry (FSCV). In section II we describe the overall concept and design of the dual-mode chip. Next, details of the circuitry used in the electrochemical amplifier as well as its performance will be discussed in section III. The design and function of the electrophysiology amplifier are presented in section IV. In section V, information about the dual-mode CMOS chip’s layout and fabrication are provided. Finally, conclusions and discussions related to the dual-mode chip will be given in section VI.

**Fig. 1.**
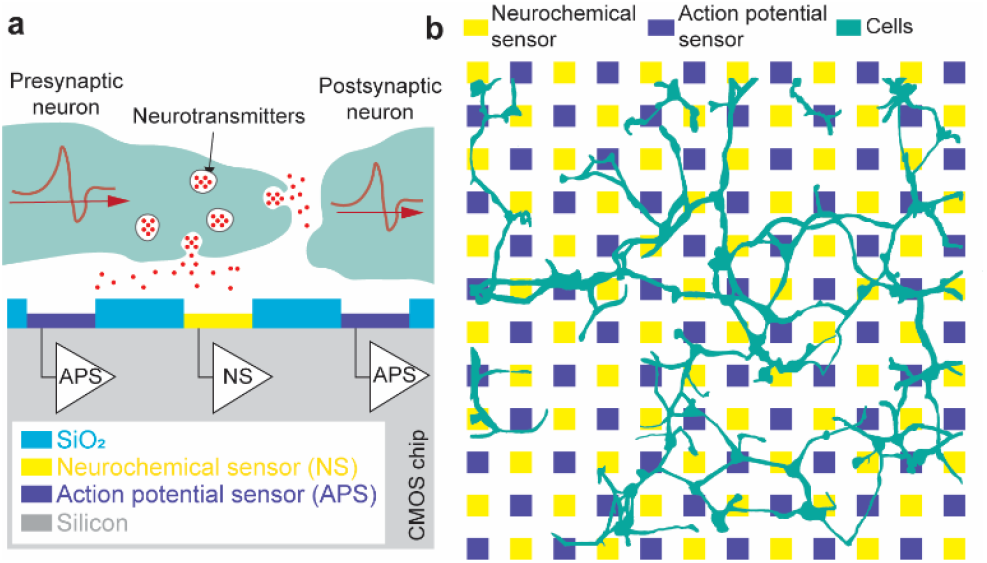
Conceptual view of the dual-mode device. (a) Cross-section showing simultaneous measurement of presynaptic action potential, neurotransmitter release at synapse, and postsynaptic action potential using dual-mode CMOS chip. (b) Top-down view showing large network of neurons located on the dual-mode microelectrode array.

## II. Dual-mode Chip Concept

The dual-mode concept is to simultaneously record neurotransmitter release and action potentials with a high electrode count, and high-density array. To create a large array of amplifiers, their design must be compact and scalable. This section describes simplified models of the electrochemical and electrophysiology amplifiers used in the dual-mode chip to achieve the goals of measurement of both signal modalities while enabling a high electrode count without sacrificing performance. More detail about the actual amplifier designs used on the fabricated dual-mode chip as well as their measured performance will be provided in the subsequent sections.

### A. System design

The dual-mode chip integrates three main functional blocks: a microelectrode array, 256 TIA neurochemical amplifiers, and 256 TCA electrophysiology amplifiers (Fig. 2). Each and every electrode on the MEA is connected to its own dedicated amplifier. Using multiplexing, all 512 amplifiers are fed to class AB output buffers for simultaneous parallel measurement using an off-chip data acquisition system [24]. The chip also contains additional circuitry for controlling the time-division multiplexing scheme, programming of SRAMs for gain control and testing modes, as well as on-chip current sources for electroplating of the electrodes with a suitable electrode material such as platinum black for potential measurements.

**Fig. 2.**
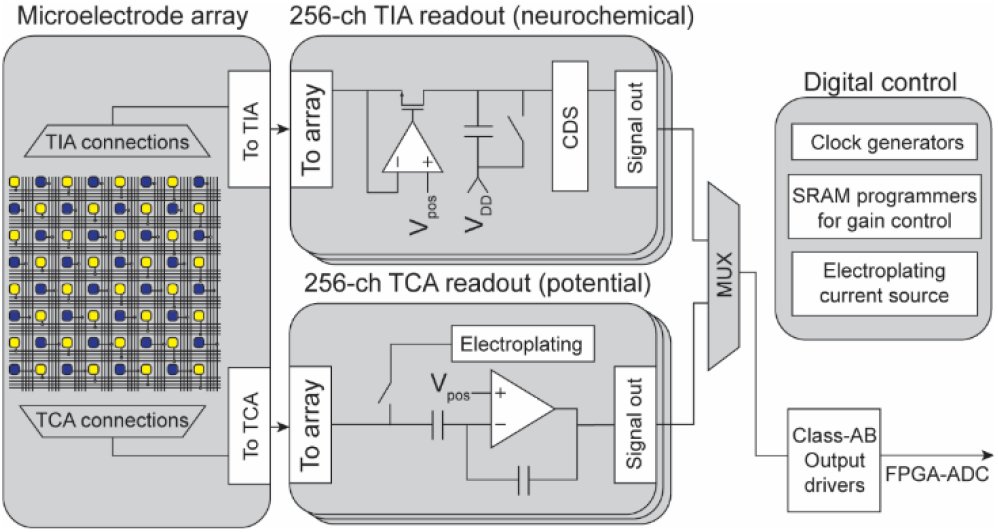
Block diagram of the dual-mode chip. Simultaneous measurement is enabled by multiplexing 256 parallel channels of neurochemical amplifiers and 256 parallel channels of electrophysiology amplifiers each of which is connected to its own electrode on a high-density microelectrode array. Additional on-chip circuitry controls the multiplexing, allows for electroplating of the electrodes, programs amplifier gain settings, and drives the off-chip ADCs.

### B. Fundamental transimpedance amplifier concept

The dual-mode chip’s electrochemical amplifier is based on our half-shared TIA design (Fig. 3a) [25]. This circuit converts electrode current by clamping the electrode voltage at a known level, V_pos_, and integrating the current onto a capacitor. Before reaching C_int_, the current must pass through the cascode transistor M_2_ and the current mirror formed by M_3_ and M_4_. The current mirror is designed so that the W/L ratio of M_3_ can be changed which causes more or less current to flow in C_int_ for the same electrode current, thus providing multiple transimpedance gain settings. The transistor M_4_ is part of a current mirror that serves two purposes. The first is to add an offset current to the measurement so that both oxidation and reduction currents can be measured by the amplifier while maintaining the same direction of current flowing through M_2_ and C_int_. The second is to enable accurate calibration of TIAs. A multiplexer (M_6_ and M_7_) is used to read out the integrated voltage. At the sample rate (*f*_*s*_), the integration capacitor is reset using M_5_ by discharging the capacitor to V_reset_. The gain can be determined by the following expression:

**Fig. 3.**
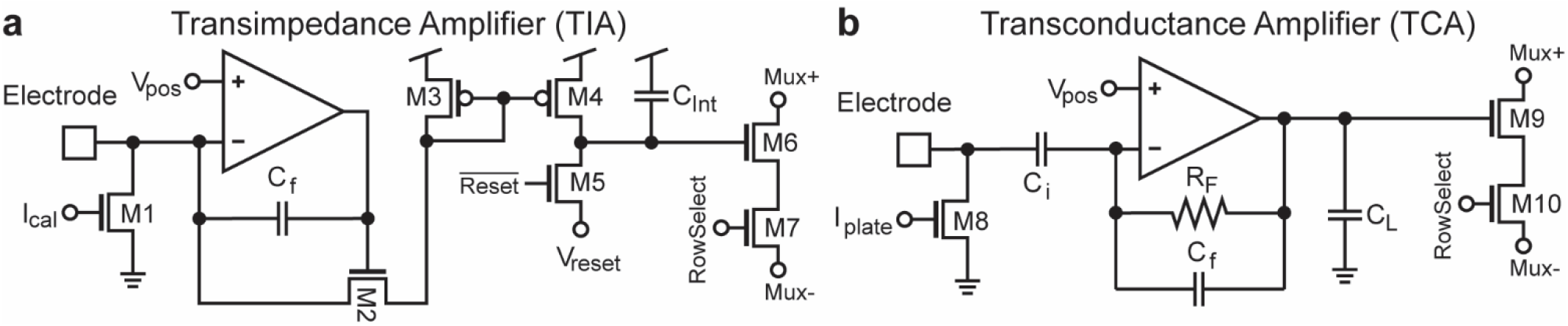
Simplified schematics of the TIA and TCA (a) The integrating TIA is based on a regulated cascode amplifier that integrates electrode current onto C_int_ through the adjustable ratioed current mirror formed by M_3_ and M_4_ to enable different gain settings. (b) The TCA is a capacitive feedback design to measure electrode potential. The output current is passed to C_L_ to produce a voltage. Both the TIA and TCA include output multiplexing circuitry and current mirrors on the electrodes to enable electroplating of suitable electrode materials such as gold, platinum black, or Ag/AgCl.

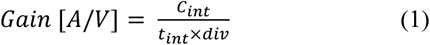

where *C*_*int*_ is the value of the integration capacitor, *t*_*int*_ is the integration period (1/*f*_*s*_) and *div* is the current mirror divider ratio. Because of the integration and sampling, the expected frequency response will correspond to the *sinc()* function [25].

### C. Fundamental transconductance amplifier concept

The electrophysiology amplifier (Fig. 3b) is based on a neural amplifier design that is commonly used with some simplifications [26]. It uses capacitors C_i_ and C_f_ to AC couple the electrode and to set the gain of the amplifier. Resistor R_f_ is a high-value resistance to provide DC stability for the amplifier. The output of the TCA drives a load capacitor C_L_ to produce an output voltage. Some important design equations for this amplifier are given below for the gain and bandwidth:

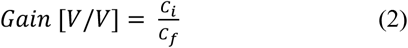

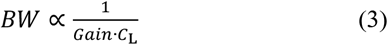

Like the TIA, an output multiplexer (M_9_ and M_10_) and an electroplating current mirror (M_8_) is included in the design.

## III. Electrochemical Amplifier

The electrochemical amplifier that is used in the dual-mode chip builds on our previously published amplifier design which is one of the highest performing transimpedance amplifier arrays published [14], [23]–[25], [27]. For the dual-mode chip, the electrochemical amplifier can operate two modes of operation, amperometry and cyclic voltammetry. Cyclic voltammetry (CV) is a method to study neurotransmitters as well as other electroactive molecules. FSCV or fast-scan cyclic voltammetry can achieve higher temporal resolution and sensitivity compared to slow-scan CV [28]–[30]. We have been able to successfully perform traditional slow-scan CV using our previous device [27] however, to implement FSCV, a wide dynamic range and flexible gain settings are added to the dual-mode chip’s electrochemical amplifiers. This section will describe the circuitry used in the electrochemical amplifiers, how FSCV is implemented, and give some performance results from the fabricated device.

### A. Amperometry mode

Amperometry mode for the electrochemical amplifier is implemented using a design that we have previously reported [14], [23]–[25], [27]. The portion of the schematic to the left of the dashed line in Fig. 4a shows the reused elements of the electrochemical sensor. Briefly, this circuit is an integrating transimpedance amplifier comprised of a half-shared op-amp [25] with a cascode transistor M_9_, where electrode current is integrated onto C_LoG_, or onto the parasitic capacitance of the node at the drain of M_9_, depending on the amperometry gain setting (changed by M_8_ and LoG). The potential of the electrode is maintained at V_pos_ due to the op-amp’s feedback through the cascode transistor. The addition of M_10_, M_12_, and M_15_ allows disconnection of the reset transistor M_4_, disconnection of the output multiplexer circuitry, and connection to the additional FSCV mode circuitry to the right of the dashed line. More information on the multiplexing operation is presented in section III.D.

**Fig. 4.**
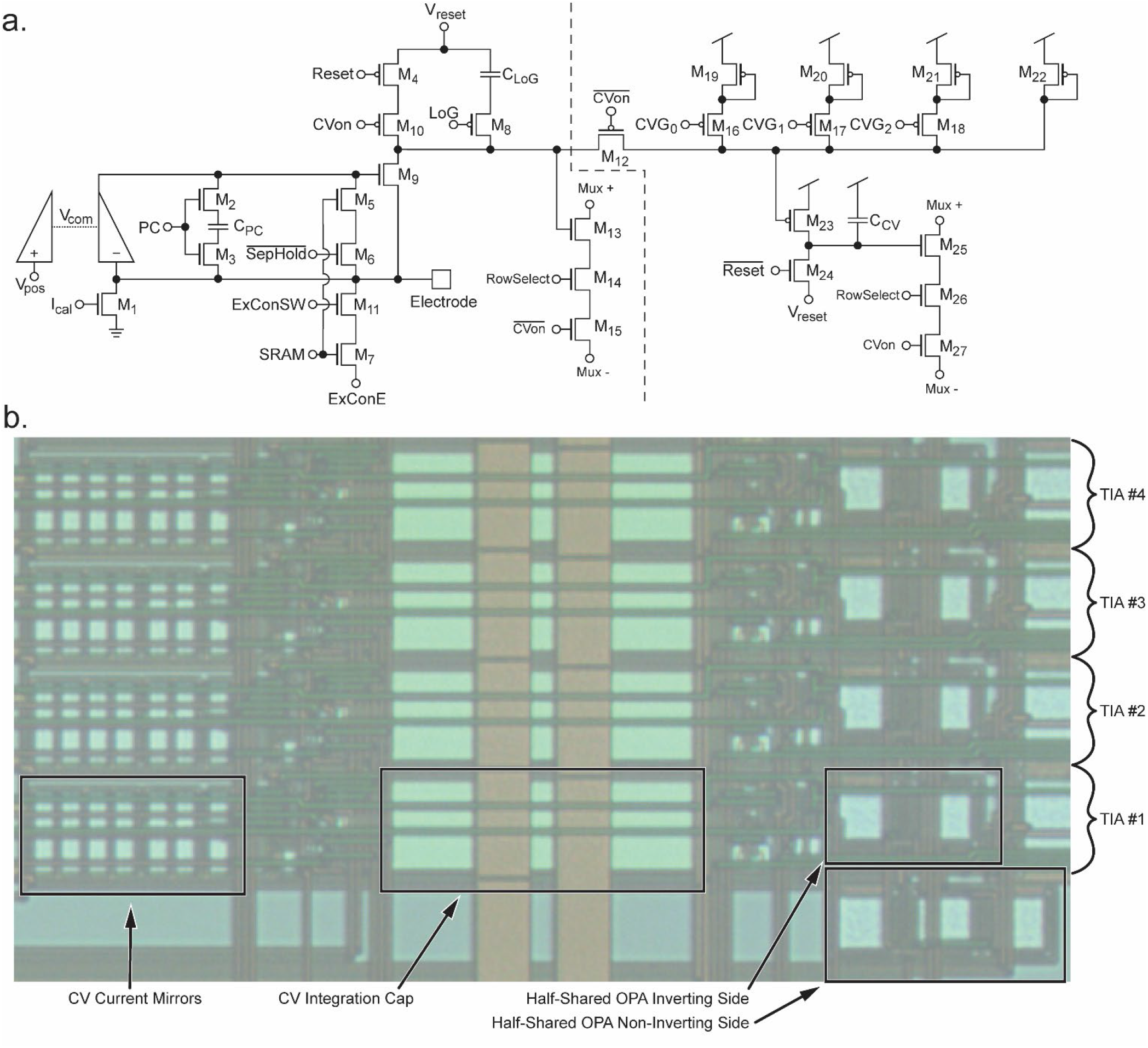
Details of the TIA’s design. (a) Schematic of the TIA. To the left of the dashed line, the schematic is similar to our previously reported TIA design and is used for amperometry. The circuitry to the right of the dashed line implements FSCV functionality using the current mirrors formed by M_19_-M_22_ and M_23_ with integration cap C_CV_. Switches M_16_-M_17_ enable multiple gain settings for FSCV mode. (b) Image of a group of 4 half-shared TIAs on the fabricated chip. Four amplifiers share the non-inverting side of the OPA to conserve space. [25] The CV integration cap and CV current mirrors occupy a considerable die area.

### B. Fast scan cyclic voltammetry mode

FSCV measurements require a wide dynamic range in the range of μA rather than pA to nA seen in amperometry, because of the large displacement current coming from the double-layer capacitance of the electrode-electrolyte interface. In the dual-mode chip, adjustable ratio current mirrors and a larger integration capacitor are included to provide a significantly higher dynamic range which corresponds to lower gain settings. These changes are shown in Fig. 4a to the right of the dashed line. M_10_ is added to enable switching to CV mode when Cv_on_ is set high. This disables the reset signal from resetting the capacitance at the drain node of M_9_. M_12_ is enabled when CV_on_ is high, connecting the original circuit to the new additional CV mode circuity. Conceptually, M_19_-M_22_ form a ratioed current mirror with M_23_ whose ratio is programmable by enabling or disabling the signals CVG_0_-CVG_2_. The current is then integrated onto the capacitor C_CV_ which is reset by M_24_. The output is sent to the output buffers through output mux transistors M_25_-M_27_. This configuration allows for the two original TIA’s gain settings set by the parasitic capacitance at the drain node of M_9_ or the integration capacitor C_LoG_, as well as new lower gain settings for FSCV. With these current mirrors and larger integration capacitor, FSCV mode allows for gain settings in the range of ∼0.48 to 1.6 V/μA which greatly extends the range into levels more suitable for FSCV, where signals typically have amplitudes in the range of a few microamperes. A photomicrograph of a group of four TIAs fabricated on-chip is shown in Fig. 4b. Four amplifiers are shown to demonstrate the space-saving half-shared structure of the op-amps along the right side of the layout. The large C_CV_ integration cap is seen in the middle of the layout. This capacitor would need to be made considerably larger to achieve the same gains if not for the next largest group of components, the current mirrors for FSCV mode, which can be seen on the left. The total size of an individual amplifier is 180.15 μm by 20.45 μm, or ∼3070 μm^2^.

### C. Electrochemical amplifier performance

The transimpedance gain of individual TIAs can be directly measured by injecting known currents into the built-in I_cal_ current mirrors (M_1_) and plotting the difference in output voltage against the input currents. For amperometry mode, current was input ranging from 0 to 9 nanoamperes, resulting in a voltage difference ranging from 0 to ∼1 volt (Fig. 5a). The measurements of the entire collection of 256 amplifiers are plotted in blue with dots for the mean and error bars showing the standard deviation. A linear fit is performed to show the gain of the TIA (red line), resulting in a transimpedance gain of 144 mV/pA at a sample rate of 40 kHz. Notice that the gain tapers off at currents above ∼7 nA, this is caused by limitations of the output buffers. Data points above this level are excluded from the linear fit.

**Fig. 5.**
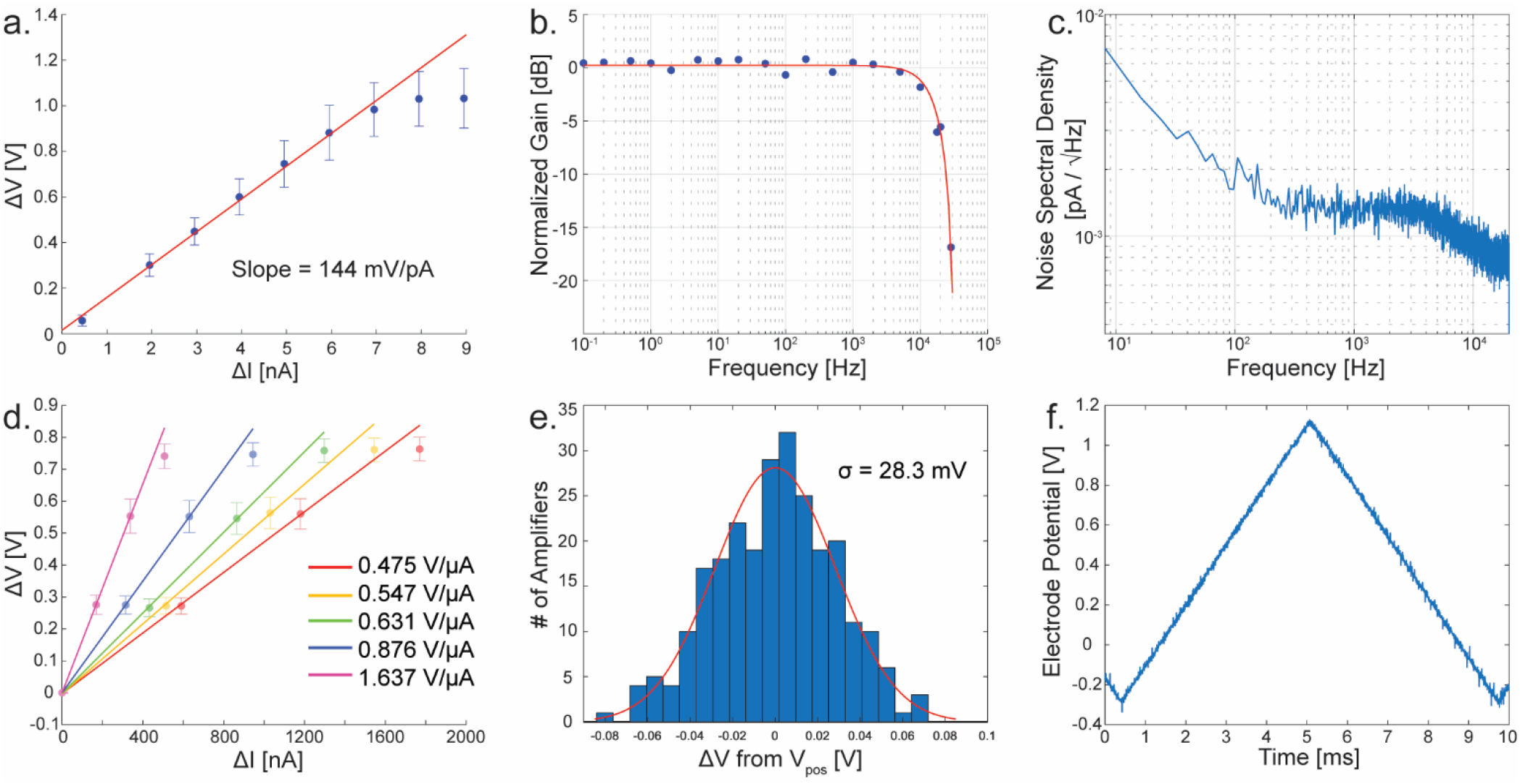
Measurements of the TIA with amperometry and FSCV modes. (a) Gain calibration in amperometry mode. (b) Frequency response at measured points (blue) with *sinc()* response best fit (red). (c) Noise spectral density of a typical TIA. (d) Gain calibration of the multiple FSCV gain settings. (e) Histogram showing the mismatch distribution between all 256 TIAs in FSCV mode. (f) Measured electrode potential of an amplifier in FSCV mode. The scan rate is 300 V/s.

The frequency domain characteristics of the TIA are also studied (Fig. 5b and c). A frequency response was obtained by enabling the external electrode connection to an amplifier and injecting a small sine wave into the electrode through a resistor to produce a known AC current. In Fig. 5b, the result of this measurement is shown. Blue points are the measurements at test frequencies ranging from 0.1 Hz to 30 kHz. The response of the integrating amplifier is known to correspond to a *sinc()* function [25], and by fitting to this response produces a cutoff frequency of ∼10.3 kHz. A noise spectral density is presented in Fig. 5c. Integrating under this curve yields a total noise of 4.51 pA_RMS_.

For FSCV mode, there are five distinct gain settings. Each of these was measured in the same way as previously mentioned for amperometry mode. In Fig. 5d these measurements are shown where each setting is plotted as its own colored points and fit line. The resulting gain settings range from ∼0.48 to 1.6 V/μA when sampled at 40 kHz. To determine the matching characteristics of the current mirrors, I_cal_ is grounded to ensure zero input current, and the resulting output voltage is subtracted from the V_pos_ level, a histogram is plotted and shown in Fig. 5e. By fitting the data of all 256 channels, the standard deviation of this mismatch is 28.3 mV. Because the measurements are calibrated to a zero current input (Fig. 5d), the voltage mismatch between channels does not harm the performance of the amplifier until the mismatch becomes so great as to limit the dynamic range of the circuit. To implement FSCV, it is necessary to stimulate the electrode potential. Typically, this is performed using a triangular-shaped pulse with a ramp rate of 300 V/s with an amplitude range up to 1.7 V [29], [30]. To ensure that the amplifier is indeed capable of driving the electrode at such speeds, a triangle wave with a ramp rate of 300 V/s and amplitude of 1.4V was applied to V_pos_ while the electrode potential was measured using ExConE. The result is shown in Fig. 5f. The amplifier shows no distortion of the triangular shape at these speeds and amplitude ranges. A comparison of the TIA to other reported designs is presented in Table I.

**TABLE I.** Comparison to Similar Transimpedance Amplifiers

| Reference | [31] | [22] | [32] | [13] | [21] | [20] | Dual-Mode TIA |
| --- | --- | --- | --- | --- | --- | --- | --- |
| Process Node | 0.5 $\mu\text{m}$ | 0.18 $\mu\text{m}$ | 0.5 $\mu\text{m}$ | 0.5 $\mu\text{m}$ | 0.13 $\mu\text{m}$ | 0.18 $\mu\text{m}$ | 0.35 $\mu\text{m}$ |
| Die Size | - | 12 × 8.9 mm | 2.5 × 2.5 mm | 3 × 3 mm | 3 × 2 mm | 5 × 2.65 mm | 5 × 5 mm |
| Electrodes | 25 | 59760 | 100 | 100 | 1024 | 200 | 512 |
| Amplifier Size | 525 $\mu\text{m}^2$ | $\geq 0.04$ mm$^2$ | 0.06 mm$^2$ | 900 $\mu\text{m}^2$ | 8000 $\mu\text{m}^2$ | 0.03 mm$^2$ | $\sim 3070$ $\mu\text{m}^2$ |
| Parallel Channels | 25 | 28 | 25 | 100 | 4 | 200 | 256 |
| Noise | $\sim 110$ fA$_{\text{RMS}}$ | 120 pA$_{\text{RMS}}$ | 7.2 pA$_{\text{RMS}}$ | $\sim 100$ fA$_{\text{RMS}}$ | 56 pA$_{\text{RMS}}$ | 480 fA$_{\text{RMS}}$ at 110 Hz filtered | 4.51 pA$_{\text{RMS}}$ |
| Bandwidth | 2 kHz | 16 kHz | 11.5 kHz | $\sim 1$ kHz | 700 Hz | 110 Hz - 10 kHz | 10.3 kHz |

### D. Multiplexing

To facilitate the integration of hundreds of amplifiers onto a single chip the outputs of the amplifiers are multiplexed together. The multiplexing circuitry in the circuit consists of M_13_-M_15_ and M_25_-M_27_ (Fig. 4a). Depending on the mode of operation (amperometry or FSCV mode) the corresponding switch is turned on (M_15_ or M_27_). When the amplifier is selected for readout, the RowSelect signal goes high. This sets up a current path between MUX+ and MUX-. In the output buffers, additional circuitry contains an op-amp structure where the M_13_ or M_25_ is effectively connected as one of the input pair devices when connected to MUX+ and MUX-. For the neurochemical amplifiers, the outputs are also passed through correlated double sampling circuitry (CDS.) In this design, 16 amplifiers are multiplexed together. The resulting output waveforms are shown in Fig. 6. The multiplexing is digitally controlled using on-chip circuitry. One of the important signals is TimingD, which is shown in Fig. 6a. This signal has a period of 40 kHz (equivalent to the sample rate.) When TimingD goes low, the multiplexing logic is reset, so that the readout of the 16 amplifiers returns to the first amplifier. Output waveforms are shown in Fig. 6b and c. The y-axis is the voltage difference between V_clamp_ and the amount of voltage integrated on the capacitor. In Fig. 6b, the amplifier is in FSCV mode, so the amount of charge on the capacitor is subtracted from V_clamp_. The output is sampled at the points shown by the red dots. This point is centered on the negative pulse produced by the CDS and is sampled at a rate that is 16× faster than the sample rate (640 kHz for a 40 kHz sample rate.) In Fig. 6c, the amplifier is in amperometry mode, and therefore the amount of charge on the capacitor is added to V_clamp_. In this case, although the samples happen at the same time with respect to the TimingD pulse, the samples are now on the positive side of the output pulse.

**Fig. 6.**
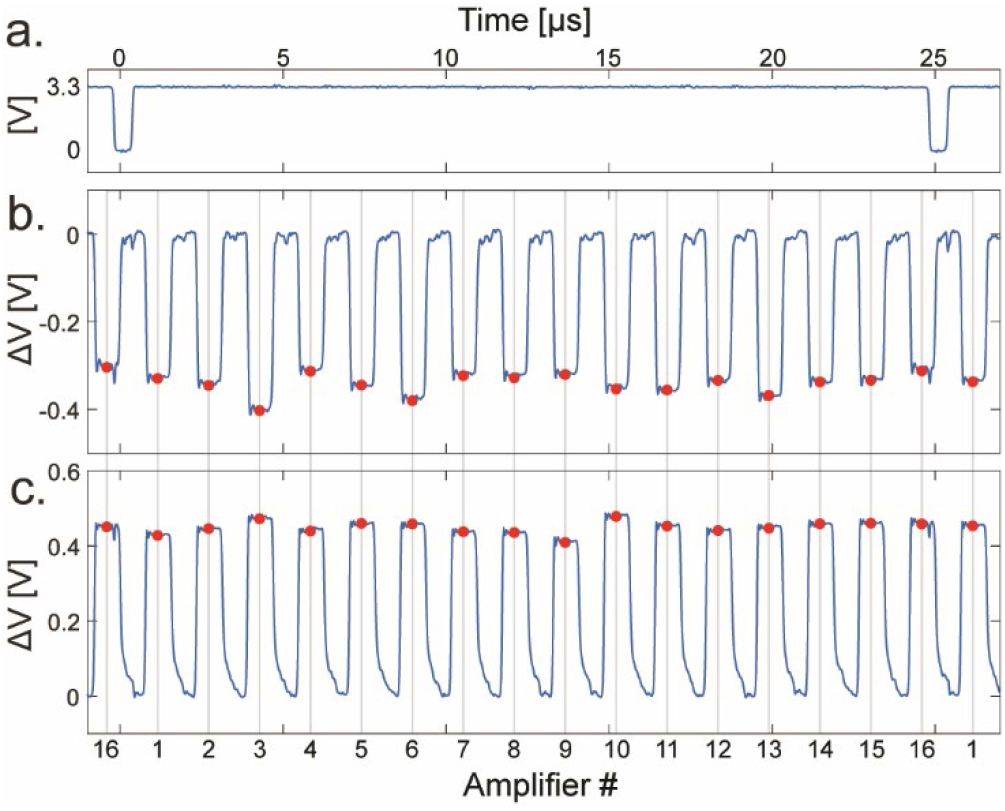
Detail of multiplexing scheme. (a) The TimingD signal, resetting of the multiplexers during the low period. (b) Signal from output buffer in FSCV mode. The red dots are the points where the output is sampled. (c) In amperometry mode, the polarity is reversed, and the signal is sampled on the peaks of the output waveform.

## IV. Electrophysiology amplifier

In order to create a dense amplifier array, the design and performance of an electrophysiology amplifier must be compact while still providing low noise and high performance to be effective at measuring neuronal signals. In on-CMOS electrophysiology, measurement of action potential and local field potentials is often achieved using transconductance amplifiers which are usually based on a design originally published in [26]. This design also provides the basis for the electrophysiology amplifier used in the dual-mode chip. This section will describe the circuit design and modifications to the common design, as well as the performance of amplifiers on the fabricated dual-mode chip.

### A. Transconductance amplifier design

The circuit used in the dual-mode chip to implement the TCA is based on a design widely used [26], however, miniaturization has been made to facilitate high-density integration and scalability. The circuit topology of the miniaturized TCA design is shown in Fig. 7. Unlike the common design, the noninverting feedback capacitors are removed to greatly reduce the required die area consumption. To allow this modification, the noninverting input of the TCA is tied to a stable reference voltage known as V_pos_. The TCA stabilizes its inverting input by negative feedback through R_F_, which is implemented as a MOS-bipolar pseudoresistor (Fig. 7a). Perturbations in the voltage at the electrode are amplified by the TCA at a ratio defined by the ratio of C_i_/C_f_, including the parasitic capacitance. The TCA’s output current is converted to voltage by feeding the output to the load capacitance C_L_ which is implemented as a MOS capacitor (M_L_) to further minimize the required die area (Fig. 7b). Some ancillary circuitry is also contained in each TCA for the output multiplexing and electroplating of the electrode surface. M_plate_ forms half of a current mirror which enables electroplating when current is forced into I_plate_ and the global current mirror transistor. M_out_ acts as an output buffer when RowSelect is high and the switch M_sel_ is activated similarly to what is described in section III.D. This allows for a current path to flow between Mux+ and Mux– which are connected to the output multiplexer.

**Fig. 7.**
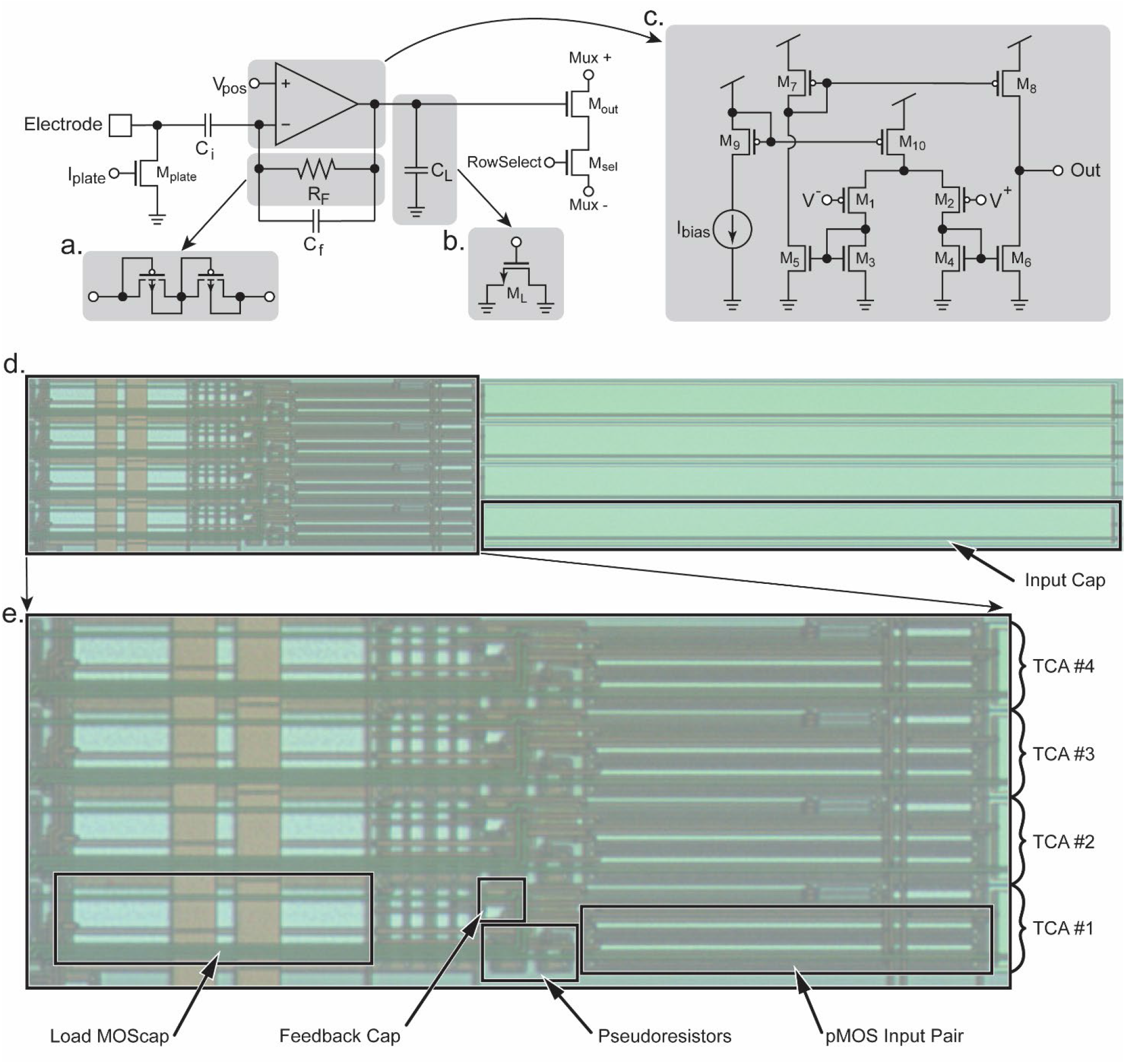
Details of the TCA’s design. The TCA uses capacitive feedback providing a gain of roughly C_i_/C_f_. (a) DC stability is provided by a pseudoresistor R_F_ which is implemented as a pair of diode-connected pMOS devices to conserve die area. (b) The load capacitor C_L_ is implemented as a MOScap for compactness. (c) The internal schematic of the multiple current mirror TCA. (d) Layout image of a group of four TCAs as fabricated. The input capacitors consume ∼60% of the die area. (e) Zoomed-in image of the active area of the TCAs. The largest components are the pMOS input devices M_1_ and M_2_ to minimize their noise contribution. The MOScap C_L_ also consumes a significant area. Comparatively, the feedback cap C_f_ and the pseudoresistors R_F_ are relatively small.

The TCA design is scaled down dramatically and adapted for the 0.35 μm process while maintaining the ideal operating regimes of the transistors (Fig. 7c) [33]. The capacitors dominate area consumption even more than the transistors. Careful consideration was taken to reduce the needed area while still yielding an effective gain, bandwidth, and noise performance of the TCA. Overall, the reduction in die area using these optimizations results in dimensions of 541.9 × 20.45 μm and a total area of 0.011 mm^2^ per amplifier which is one of the smallest reported to date. A photomicrograph of a group of fabricated TCAs is shown in Fig. 7d. Most of the area, approximately 60%, is consumed by the ∼10 pF input capacitor, emphasizing the importance of capacitor dimension optimization. Because of the large size of the input capacitor (Fig. 7e) is also presented, which shows only the active area of the TCAs. The large input pair, M_1_ and M_2_ can be seen with their extreme *W/L* ratio enabling their operation in moderate-to-weak inversion for better noise performance. The load capacitor C_L_ is the next largest component in terms of die area. Also shown are the relatively small pseudoresistors and feedback capacitor. All other components consume negligible area compared to those already mentioned.

### B. Transconductance amplifier performance

The gain and bandwidth of each TCA are obtained by injecting a small sine wave into the ExConE connection and recording the output amplitude. The response of a typical amplifier is presented in Fig. 8a. The passband gain is 37.5 dB, about 75 V/V. The bandwidth is from ∼0.2 Hz to ∼10 kHz which is sufficient to measure signals in both the local field potential (LFP) band (<300Hz) and in the action potential (AP) band (300 Hz -7.5 kHz) [34], [35]. Since APs are very low amplitude signals, the noise performance of the amplifier must be low. Fig. 8b shows a measured noise spectral density of a TCA. Integrating this noise in the AP band yields an input-referred noise of 24.9 μV_RMS_. To show the transient performance of the TCA, a measurement of a synthetic neural spike is performed at two different amplitudes (∼1 mV and 4 mV, peak-to-peak). The spikes are generated using a common scheme [36] and fed into the amplifier. Fig. 8c shows the resulting signal of a synthetic neural spike with a peak-to-peak amplitude of ∼1mV. The signal is also shown filtered to the AP band, and a closer view of a single spike is also shown. Similarly, another measurement was performed with spikes of ∼4 mV amplitude (Fig. 8d). These measurements show that neural spikes of approximately 1mV or more are easily discerned from the random spikes occurring in the noise, and the amplifiers would therefore be capable of measuring actual spikes from live neurons. Table II is shown to compare the TCA’s performance with other presented designs.

**Fig. 8.**
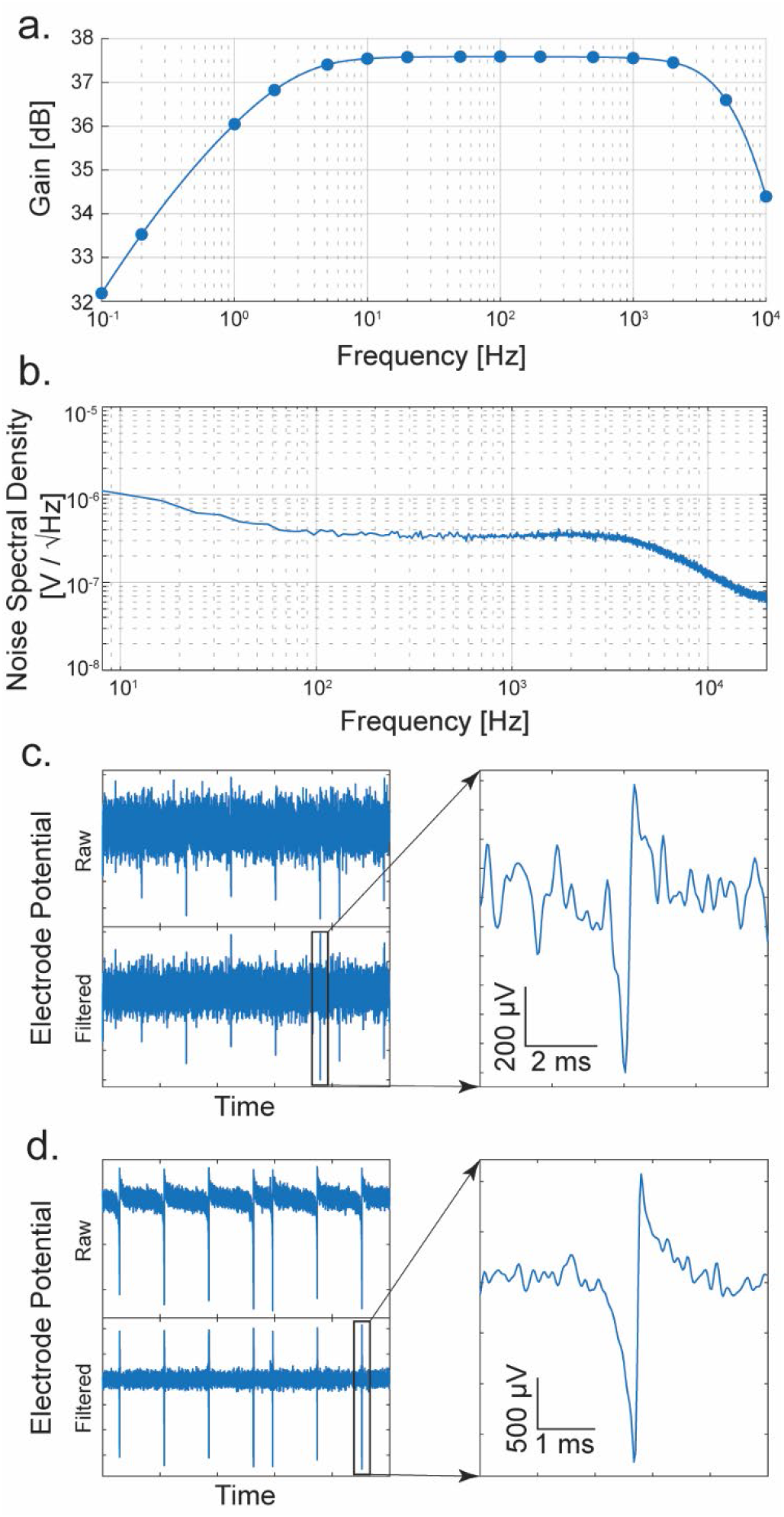
Performance of the TCA. (a) Frequency response of a typical amplifier. (b) Noise spectral density of a typical amplifier. (c) On-chip measurement of a synthesized neural spike of ∼1mV amplitude. The raw and filtered from 300 to 7000 Hz are shown, as well as a close-up of a selected spike. (d) Measurement of a ∼4mV amplitude synthetic neural

**TABLE II.** Comparison to Similar transconductance amplifiers

| Reference | [26] | [22] | [21] | [37] | [38] | Dual Mode TCA |
| --- | --- | --- | --- | --- | --- | --- |
| Noise | $2.2 \mu\text{V}_{\text{rms}}$ | $5.4 \mu\text{V}_{\text{rms}}$ | $7.7 \mu\text{V}_{\text{rms}}$ | $2.2 \mu\text{V}_{\text{rms}}$ | $6.4 \mu\text{V}_{\text{rms}}$ | $24.9 \mu\text{V}_{\text{rms}}$ |
| Sample Rate | - | 20 kHz | - | 25 kHz | 30 kHz | 40 kHz |
| Bandwidth | 7.5 kHz | 300 Hz - 10 kHz | 300 Hz - 6 kHz | 300 Hz - 10 kHz | 10 kHz | 0.2 Hz - 10 kHz |
| Size (per amplifier) | $0.16 \text{ mm}^2$ | - | - | $0.07 \text{ mm}^2$ | - | $0.011 \text{ mm}^2$ |
| Process Node | $1.5 \mu\text{m}$ | $0.18 \mu\text{m}$ | $0.13 \mu\text{m}$ | $0.35 \mu\text{m}$ | $0.13 \mu\text{m}$ | $0.35 \mu\text{m}$ |
| Voltage | 5 V | - | - | - | 1.8 V | 3.3 V |
| Current | $8 \mu\text{A}$ | - | $2.5\text{-}37.5 \mu\text{A}$ | - | - | $8 \mu\text{A}$ |
| Power | $40 \mu\text{W}$ | $16 \mu\text{W}$ | - | $30 \mu\text{W}$ | $49.1 \mu\text{W}$ | $26 \mu\text{W}$ |
| Parallel Channels | 6 | 2048 | 1024 | 36 | 384 | 256 |
| Electrodes | - | 59760 | 1024 | 36 | 966 | 512 |
| Electrode Pitch | - | $13.5 \mu\text{m}$ | $58 \mu\text{m}$ | $25 \mu\text{m}$ | $\sim 70 \mu\text{m}$ | $16 \mu\text{m}$ |
| Die Size | $2.2 \times 2.2 \text{ mm}$ | $12 \times 8.9 \text{ mm}$ | $3 \times 1.85 \text{ mm}$ | - | $9 \times 5 \text{ mm}$ | $5 \times 5 \text{ mm}$ |

## V. CMOS Implementation

The dual-mode CMOS chip is designed and fabricated using a typical 0.35 µm 4 metal 2 poly N-well CMOS process. For this initial concept device, a die size of 5×5 mm is used (Fig. 9), resulting in 512 fully parallel channels, with 256 dedicated to electrochemical measurements and 256 to electrophysiology measurements. This section describes considerations made in the layout of the device, as well as details of the on-chip microelectrode array.

**Fig. 9.**
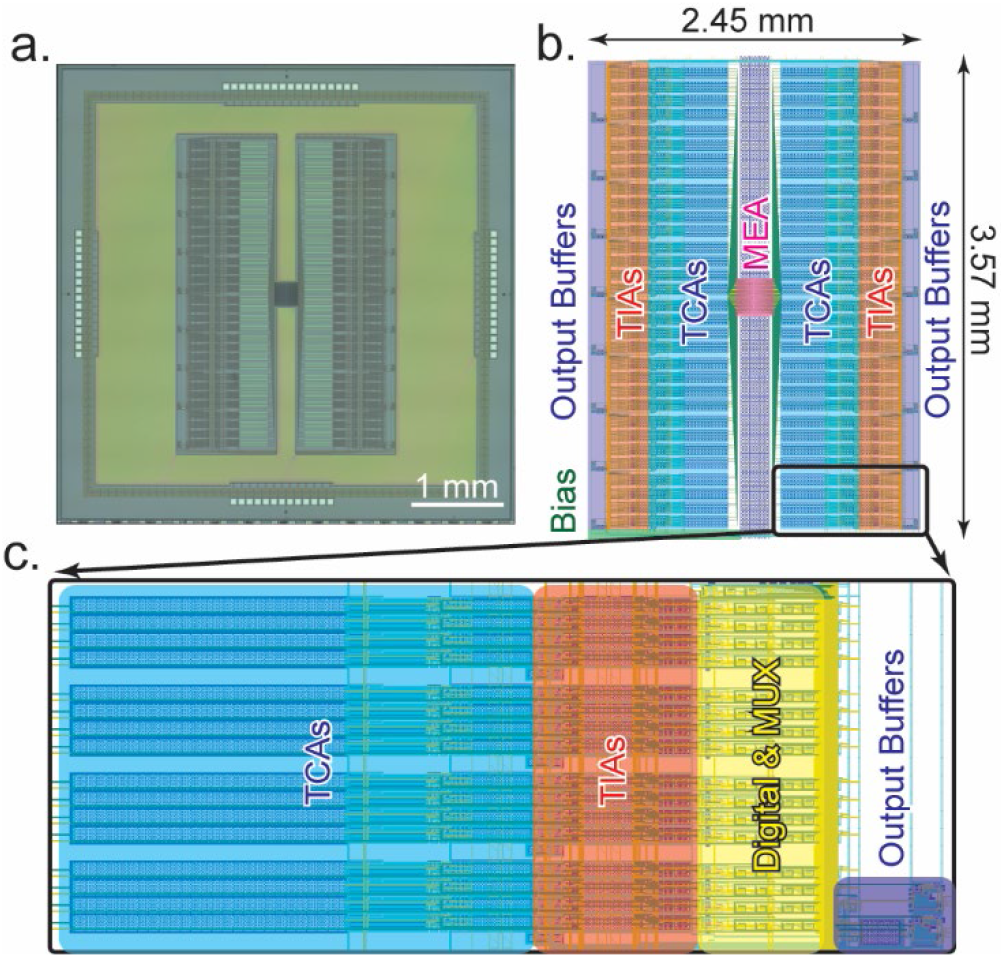
Layout information of the dual-mode chip. (a) Photomicrograph of the fabricated dual-mode chip using 0.35 µm CMOS technology. The total size is 5×5 mm. (b) Location of various structures on the chip. Amplifiers are mirrored on both sides of the centrally located microelectrode array. The active area is 2.45×3.57mm. (c) Close-up view of a “pseudocolumn” showing how each group of 16 amplifiers is arranged to enable multiplexing into a single output buffer.

### A. Chip layout

The microelectrode array is placed at the center of the die (Fig. 9a). In Fig. 9b an annotated view of the layout’s active area is shown. Wiring from the electrodes fans out to both sides of the array to connect to the amplifiers. On the left and right sides of the MEA, the amplifiers, digital circuitry, output multiplexing, output buffers, and biasing circuitry are mirrored. This arrangement gives a compact use of space and simplifies the electrode interconnection wiring. The dual-mode chip is designed to be interfaced with a custom-designed data acquisition system, so the 512 amplifier channels are multiplexed into 32 output channels [24]. To accomplish this, the outputs are grouped into 16 current channels and 16 voltage channels. These are then split into 8 and 8 on each side of the chip. Thus, each dedicated output serves groups of 16 amplifiers. In the close-up section of Fig. 9c, this grouping of 16 amplifiers can be seen. Each of these groups of 16 contains 16 TIAs, 16 TCAs, and 2 output buffers. They are multiplexed together using digital control circuitry and the signals are buffered by the output buffers which are then connected to wire bonding pads to drive the off-chip ADCs. Within the group of 16 amplifiers, the TIAs are grouped in groups of 4 for half sharing. This is clearly seen in both Fig. 9c and Fig. 4b with the gaps between the groups of 4 amplifiers. For future designs, further space optimization could be obtained, for example, by rearranging the TCA input capacitors to use this empty space. For this initial design, this was not done to simplify the layout and ensure that all amplifiers are as identical as possible to minimize any potential peculiarities caused by nonidentical layout.

### B. Microelectrode array

The microelectrode array is a critical part of the dual-mode chip since it is the source of the signals that we are aiming to measure. It must have a high enough spatial resolution to resolve fine details between neurons. To determine a reasonable pitch, we examined other literature and chose a value on a similar scale. In other MEAs some typical ranges are from 13.5 μm [22], 18 μm [39], to 42 μm [19] and even wider. For this design, the value was chosen to be 16 μm between electrodes. This value was chosen to stay closer to the smallest pitch reported in the literature, while giving sufficient space for the routing of the wiring out from the electrodes to the amplifiers as well as potential post processing that my take place. The resulting electrode array is shown in Fig. 10. In Fig. 10a, the layout is shown. The routing of the wiring fanning out from the center to the left and right sides, where they run out to connect to the amplifiers. The overall size of the MEA is 255.4 μm on each side. A closer view is shown in Fig. 10b. The opening in the top glass passivation layer is 5 μm which is seen as an orange-colored square in the opening. This is the exposed area of AlCu where signals will enter the chip. Between similar mode electrodes, the spacing is 16 μm and between differing mode electrodes, the pitch is 8 μm. The array is effectively two separate arrays with a pitch of 16 μm between adjacent electrodes. One for the voltage channels and one for the current channels. Each array is interdigitated at an 8 μm pitch to produce a checkerboard pattern that can be seen in Fig. 10c. Finally, a fabricated MEA is shown in Fig. 10d. The density of the MEA is close to the limit of what can practically be fabricated in 0.35 µm technology without deliberately violating design rules which would potentially result in shorted electrodes or non-exposed electrodes on the chip’s surface. For effectiveness in practical measurements, the electroplating circuitry within the amplifiers can be used to deposit a suitable and biocompatible electrode surface such as gold, platinum black, or Ag/AgCl.

**Fig. 10.**
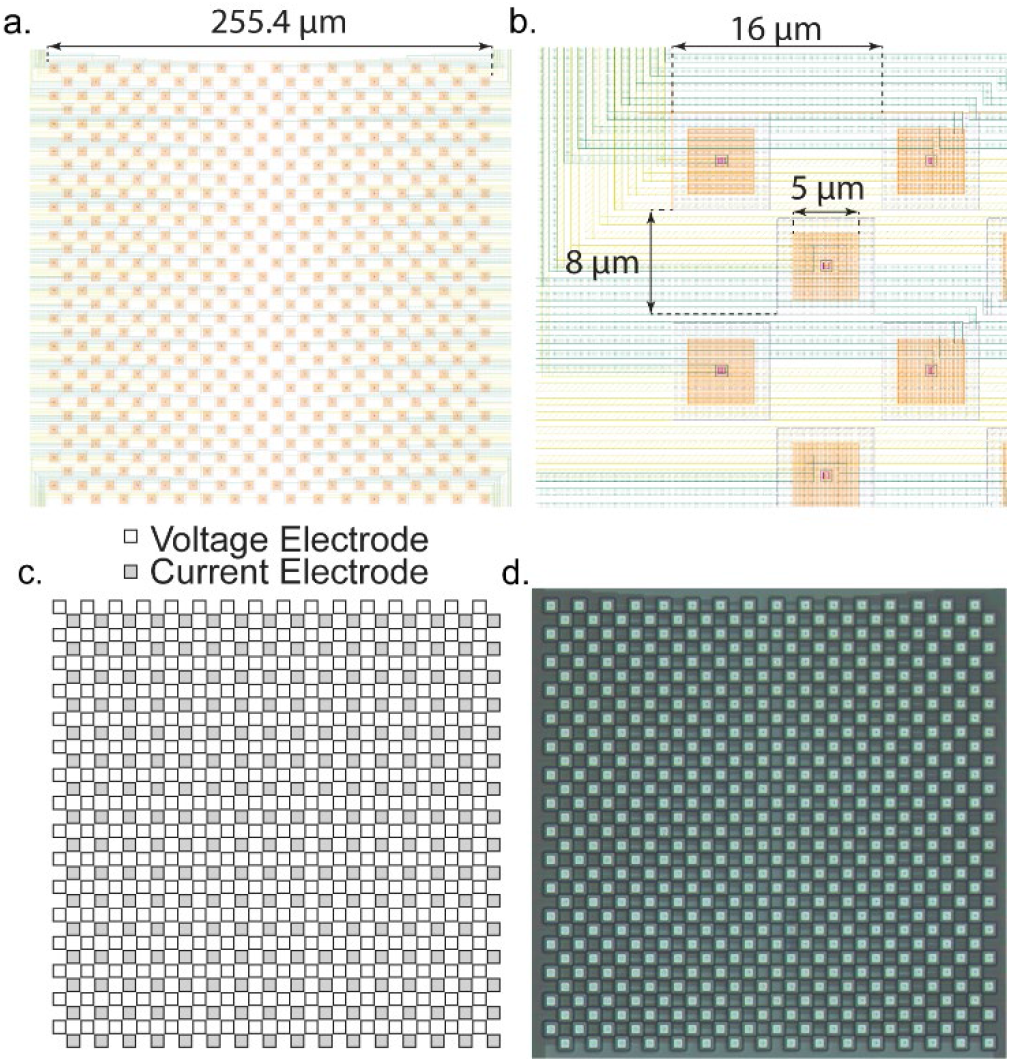
Design of the microelectrode array. (a) Layout image of the MEA. The MEA is 512 electrodes with 256 dedicated to voltage measurement and 256 dedicated to current measurement. The total size is ∼250×250 µm. (b) Detail of the individual electrodes. The pitch between electrodes is 16 µm, the glass opening is 5 µm, and between rows the pitch is 8 µm. Each row is staggered, forming the checkerboard pattern. The yellow and green layers are the metal interconnects that connect the electrodes to the amplifiers. (c) Arrangement of the voltage and current electrodes. Each row serves a different modality amplifier. (d) Photomicrograph of the MEA as fabricated on-chip.

## VI. Conclusion and Discussion

We present a CMOS chip that is capable of a fully parallel and simultaneous measurement of 512 channels from an on-chip microelectrode array with 256 neurochemical and 256 electrophysiology amplifiers. Since the amplifiers are compact and scalable, a larger array with 1000s of amplifiers could easily be integrated into a chip that would fit within a commonly used silicon reticle. This proof-of-concept design fits 512 amplifiers and an electrode array into a total active area of only 2.45×3.57mm. This dual-mode technology allows the simultaneous study of the interactions between neurotransmission, synaptic function, and action potential propagation. As a tool, this technology could provide a means for neuroscientists to study the mechanisms of neuronal degradation and synaptic dysfunction seen in cases of many neurodegenerative disorders.

For future implementations, an increase in electrode count would be desired to enable measurements of larger physical areas. Additionally, a potential improvement to this design applies specifically to the design of MEA itself. The soma of a neuron is roughly equivalent in size to our MEA’s electrode pitch, but a neuron will typically have multiple dendrites for receiving neurotransmitters from presynaptic neurons. This fact would suggest that if a more complete analysis of the interactions between neurotransmission and action potentials is desired, then an optimal MEA would have more neurochemical electrodes than electrophysiology electrodes. However, the dual-mode device in its current form is still the first device reported that is capable of measuring neurochemical signals at a similar spatial resolution as action potentials, and the design of a dual-mode chip with an improved MEA would be simple as the amplifier designs are compact and scalable.

